# Phasic dopamine drives conditioned responding beyond its role in learning

**DOI:** 10.64898/2026.03.25.714259

**Authors:** Jay A. Hennig, Mark Burrell, Naoshige Uchida, Samuel J. Gershman

## Abstract

Animals exposed to pairings of a neutral stimulus with reward acquire a conditioned response to the neutral stimulus. A prominent hypothesis, formalized in the Temporal Difference (TD) learning algorithm, is that animals learn to predict the future reward associated with the neutral stimulus (“value”). Though the TD algorithm does not explicitly specify what drives conditioned responding, a typical assumption is that it reflects the animal’s estimate of value. In TD learning, value estimates are updated using reward prediction error (RPE, the discrepancy between observed and predicted reward), and are thought to be signaled by the phasic activity of midbrain dopamine neurons. This hypothesis posits that dopamine’s effects on conditioned responding are mediated entirely by its effects on learning. However, recent experimental and theoretical evidence suggests that dopamine may play a more direct role in modulating conditioned responding. We use a combination of data analysis and computational modeling to probe the relationship between dopamine and conditioned responding. Our results suggest that dopamine directly modulates conditioned responding, in addition to its role in learning. These findings can be captured by a model in which dopamine RPE acts both indirectly (via learning) and directly on conditioned responding.

## Introduction

In Pavlovian conditioning, animals learn to associate a neutral conditioned stimulus (CS) with the delivery of an appetitive or aversive stimulus (US). For example, when a neutral odor or tone cue is repeatedly paired with a subsequent water reward, rodents exhibit anticipatory licking of the water spout during cue presentation. Pavlovian conditioning is often framed through the lens of reinforcement learning theory, which posits that conditioned responding reflects a learned estimate of the cumulative future reward (*value*) following the conditioned stimulus [1, 2, 3]. A leading theory of how animals acquire value estimates is the Temporal Difference (TD) learning algorithm, which updates value estimates using a reward prediction error (RPE), the discrepancy between observed and predicted reward. The RPE is thought to be signaled by the phasic spiking activity of dopamine neurons in the midbrain [4, 5, 6, 7]. This hypothesis is supported by a quantitative match between RPEs and dopamine neuron activity [8, 9, 10, 11], as well as by perturbation experiments establishing the causal role of dopamine in Pavlovian conditioning [12, 13, 14, 15].

The dopamine RPE hypothesis holds that dopamine’s role in Pavlovian conditioning is delimited entirely by its role in learning. Any effects dopamine has on conditioned responding must, according to this account, be indirect and delayed. We will challenge this account, presenting data showing that dopamine has a direct, immediate effect on conditioned responding. Several studies already point in this direction [16, 17, 18], though the evidence is mixed [15]. A direct role has been posited by some computational models [19, 20, 21, 22, 23], and might be mediated at the cellular level by the effect of dopamine on the excitability of striatal neurons [24, 25]. However, a systematic empirical investigation of this hypothesis has yet to be undertaken.

Disentangling the direct and indirect effects of dopamine on conditioned responding is intrinsically challenging because they cannot be measured separately, and they are often correlated. We address these challenges by conducting a fine-grained analysis of the trial-by-trial covariation between dopamine neuron activity and conditioned responding. Using computational models, we show that the patterns of covariation are most consistent with a model in which dopamine acts both indirectly via learning and directly via modulation of conditioned responding. This model can also account for the heterogeneous effects of optogenetic perturbations on conditioned responding. Taken together, our findings demonstrate that dopamine’s role in Pavlovian conditioning goes beyond learning, inviting a reconsideration of models positing a direct role in response generation.

## Results

### Conditioned responding during contingency degradation covaries with trial-averaged dopamine, and not value

We start by revisiting TD learning in the context of trace conditioning (see Methods for mathematical details). In each trace conditioning trial, the animal is presented with a cue (“CS”; e.g., a particular odor), followed by a delay, and then a reward (“US”; e.g., a drop of water; Fig 1A). In a typical TD learning model of trace conditioning, animals learn to estimate the moment-by-moment *value*, or the expected discounted cumulative reward, using RPEs (Fig 1B). The phasic responses of midbrain dopamine neurons to both the CS and US are well described by the RPE signal (see [7] for a review), supporting their hypothesized role in TD learning and the interpretation of dopamine activity as a putative measure of the animal’s RPE.

**Fig 1.**
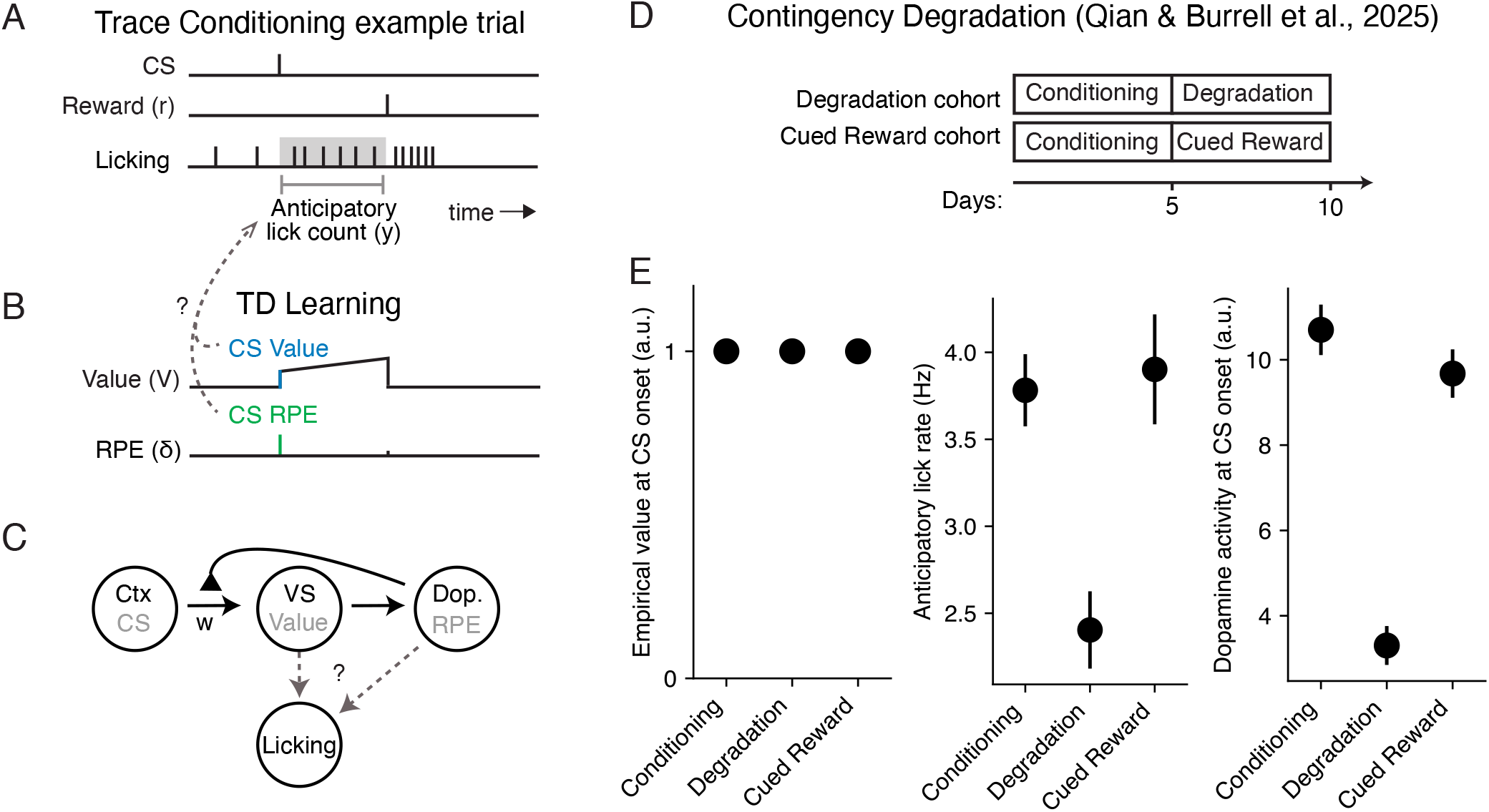
TD learning and conditioned responding during trace conditioning with contingency degradation. **A**. Example trace conditioning trial consisting of a conditioned stimulus (CS), followed by a delay, and then a reward stimulus. Also depicted is an example of the times the animal licked the reward spout, with anticipatory licking (the conditioned response) measured during the interval between the CS onset and reward delivery. **B**. Example of the estimated value and reward prediction error (RPE) of TD learning during the example trial shown in panel A. Dashed arrows depict different potential relationships between CS Value, CS RPE, and anticipatory licking. **C**. Putative circuit implementation of TD learning, where sensory cortex provides the CS input, ventral striatum (VS) carries the value estimate, and midbrain dopamine neurons provide the RPE signal, which modulates feedforward plasticity between cortex and VS. **D**. Experimental structure of the contingency degradation experiment [26]. Two cohorts underwent a “Conditioning” phase for five days/sessions, followed by five days of either a “Degradation” or “Cued Reward” phase for the Degradation and Cued Reward cohorts, respectively. **E**. Average objective value at CS onset, anticipatory lick rate, and dopamine activity (z-scored axonal calcium fluorescence) at CS onset, for the Odor A cue during each of three phases shown in panel D. Black circles and lines denote mean *±* SE across trials.

Mice trained on trace conditioning with an odor CS and a water reward produce anticipatory licking during the delay period—their conditioned response (Fig 1A). To link the TD model to conditioned responding, we require an auxiliary assumption about response generation (Fig 1C). Conditioned responding typically covaries with the CS value. However, as we’ll see, differences in CS RPEs across cues and across time are also correlated with conditioned response rates. This raises the possibility that conditioned responding could additionally reflect CS RPEs, putatively signaled by phasic dopamine.

Recent work using a contingency degradation paradigm supports this possibility [26]. In this study, there were three distinct experimental phases, presented to two different cohorts of mice (Fig 1D). Briefly, during the Conditioning phase, each trial consisted of a CS (“Odor A”) followed by a variable reward. Mice in the Degradation cohort experienced a Degradation phase, where Odor A trials were interleaved with trials containing uncued rewards. Mice in the Cued Reward cohort experienced a Cued Reward phase, where Odor A trials were interleaved with reward trials cued by an additional odor CS. Critically, the Odor A CS was rewarded similarly across all three phases, meaning its objective value was held constant across every phase (Fig 1E, left). Notably, animals’ anticipatory licking was decreased in the contingency phase compared to the conditioning and cued reward phases (Fig 1E, middle), even though the CS value was constant across phases. This mirrored the CS dopamine response (Fig 1E, right), and was well explained by a TD learning model where CS dopamine reflects the CS RPE. This suggests that differences in conditioned responding may be driven by CS RPEs and not CS value.

While the possibility that conditioned responding is based on CS RPE and not CS value is suggestive, the evidence presented in [26] is based on trial-averaged activity and correlations across cues or phases of trials. If the CS RPE is truly driving conditioned responding, then the correspondence between dopamine and licking should also manifest on a trial-to-trial basis for a single cue.

### Trials with larger CS dopamine have higher conditioned response rates

To look for evidence of a trial-to-trial correspondence between CS dopamine and conditioned responding, we first compared the number of anticipatory licks with the peak magnitude of dopamine activity following CS onset. We first carry out this analysis on each trial of an example session at the end of conditioning for one mouse, and then show the results aggregated across all sessions and mice. For this and all following analyses of data from [26], we only consider trials with Odor A, which is the CS that was identically rewarded throughout the experiment (i.e., the objective value of this CS was constant). “Dopamine activity” here means the z-scored axonal calcium fluorescence (see Methods). We labeled trials where the peak CS dopamine activity was in the highest quartile across trials as a “High CS dopamine” trial (Fig 2A), and those where the peak in CS dopamine activity was in the lowest quartile as a “Low CS dopamine” trial (Fig 2B). In this example session (Fig 2C), trials with high CS dopamine had more anticipatory licks on average than trials with low CS dopamine (Fig 2C; one-sided Wilcoxon rank sum test, *W* = 2.638, *p* = 0.004).

**Fig 2.**
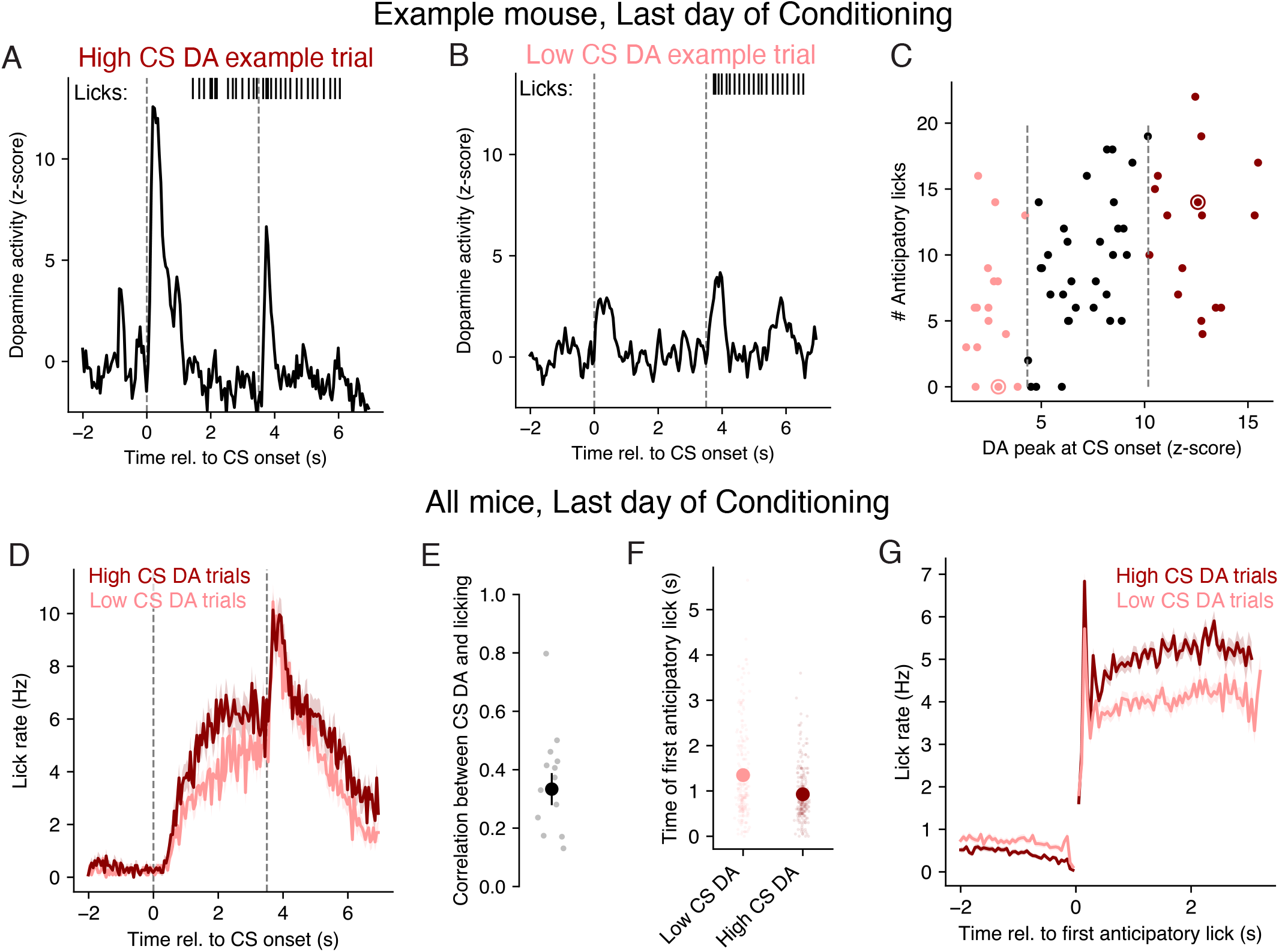
Anticipatory lick rates late in conditioning are larger on trials with larger CS-evoked dopamine. **A-B**. Dopamine activity (black trace) and licks (black vertical lines) during an example “High CS dopamine” (panel A) and “Low CS dopamine” (panel B) trial, on the last day of conditioning from an example mouse. **C**. Number of anticipatory licks and dopamine response to the CS on trials from the same example session as panels A and B. Dashed lines depict lower and upper quartiles of the CS dopamine response, used to define “Low CS dopamine” (light red) and “High CS dopamine” (dark red) trials, respectively. Dark red and light red circles indicate the example trials shown in panels A and B, respectively. **D**. Average lick rate (mean *±*SE) across all High CS dopamine and Low CS dopamine trials from the last day of conditioning in all mice. **E**. Pearson correlation between anticipatory lick rate and CS dopamine across trials in the last day of conditioning for each mouse (gray dots), and across mice (black circle). Black line depicts mean *±*SE. **F**. Average time of the first anticipatory lick (mean *±*SE) across the same trials as in panel D. **G**. Average lick rate (mean *±*SE) after aligning lick times within each trial to the time of the first anticipatory lick.

This same general pattern was also evident across sessions from all mice, where we observed that trials with high CS dopamine had higher anticipatory lick rates than did trials with low CS dopamine (Fig 2D; one-sided Wilcoxon rank sum test, *W* = 7.561, *p <* 1 *×* 10^*−*3^). This effect could not be explained by differences in satiety, as high CS dopamine trials had higher anticipatory lick rates than low CS dopamine trials even when excluding the last half of trials from each session (one-sided Wilcoxon rank sum test, *W* = 3.769, *p <* 1 *×* 10^*−*3^). CS dopamine and anticipatory lick rates had a graded relationship, such that higher CS dopamine was correlated with larger anticipatory lick rates on the last day of conditioning for every mouse (Fig 2D; average Pearson correlation was 0.33*±*0.05, mean *±* SE; *N* = 14 animals). While TD learning predicts that both CS dopamine and CS value should increase with learning, this correspondence between anticipatory licking and CS dopamine is unexpected on the last day of conditioning, when animals have fully learned the reward contingencies. This result is also consistent with empirical results in previous studies that found anticipatory lick rates were decreased when dopamine neurons were transiently inhibited during the ISI using optogenetics [18, 15], a result we will return to in a later section.

Differences in the average lick rates of rodents may be due to differences in licking latency, intensity, or duration [27, 28, 29, 30, 31]. We found that trials with high CS dopamine did have slightly shorter latencies to the time of the first lick than trials with low CS dopamine (Fig 2F; one-sided Wilcoxon rank-sum test, *W* = *−*4.374, *p <* 1 *×* 10^*−*3^). Anticipatory lick rates were also larger on high versus low CS dopamine trials when aligning to the time of the first lick (Fig 2G; one-sided Wilcoxon rank-sum test, *W* = 12.902, *p <* 1 *×* 10^*−*3^). Thus, trials with high CS dopamine had both shorter licking latencies and higher lick rates than trials with low CS dopamine.

We next asked whether these differences in lick rate as a function of the CS dopamine response were present in other studies that recorded both dopamine and licking in mice undergoing odor trace conditioning [10, 32, 11, 33]. While the effect size varied across studies, trials with high CS dopamine all tended to have higher rates of anticipatory licking than trials with low CS dopamine across a range of studies and dopamine signals (Fig 3). These results suggest that the magnitude of the CS dopamine response can, on average, explain variability in the subsequent amount of anticipatory licking. As we will show in a later section, this finding cannot be explained by the traditional TD learning assumption that CS value solely drives conditioned responding.

**Fig 3.**
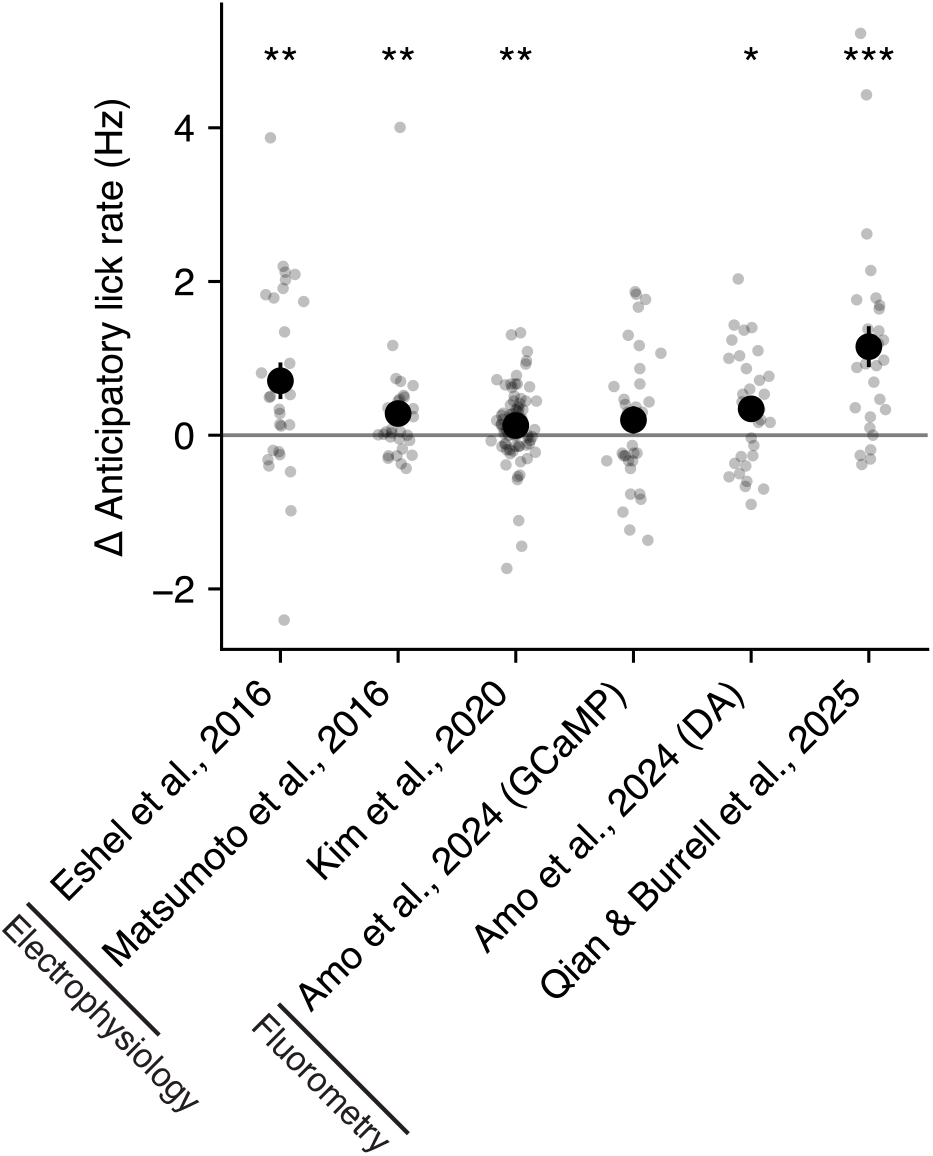
Anticipatory lick rates are larger on trials with higher CS dopamine than on trials with low CS dopamine across multiple studies. For each study, gray dots depict the average difference in the anticipatory lick rate on trials with High versus Low CS dopamine within a single session. Black circles depict the mean across sessions, and black lines depict the mean *±*SE across sessions. Asterisks depict significance of a Wilcoxon signed rank test at the 0.05 (*), 0.01 (**), and 0.001 (***) level. Black lines on x-axis tick labels indicate the dopamine signal recording technique used in that study. For [33], results are shown separately for dopamine activity measured using calcium (GCaMP) and dopamine (DA) sensors.

### CS dopamine and conditioned responding covary throughout conditioning

We next asked whether the relationship between CS dopamine and anticipatory licking observed at the end of conditioning was also present throughout the learning process. We visualized the time series of CS dopamine and anticipatory licking on each trial to the same cue during the conditioning phase of [26]. CS dopamine and anticipatory lick rates were positively correlated across all animals (Fig 4), with an average

**Fig 4.**
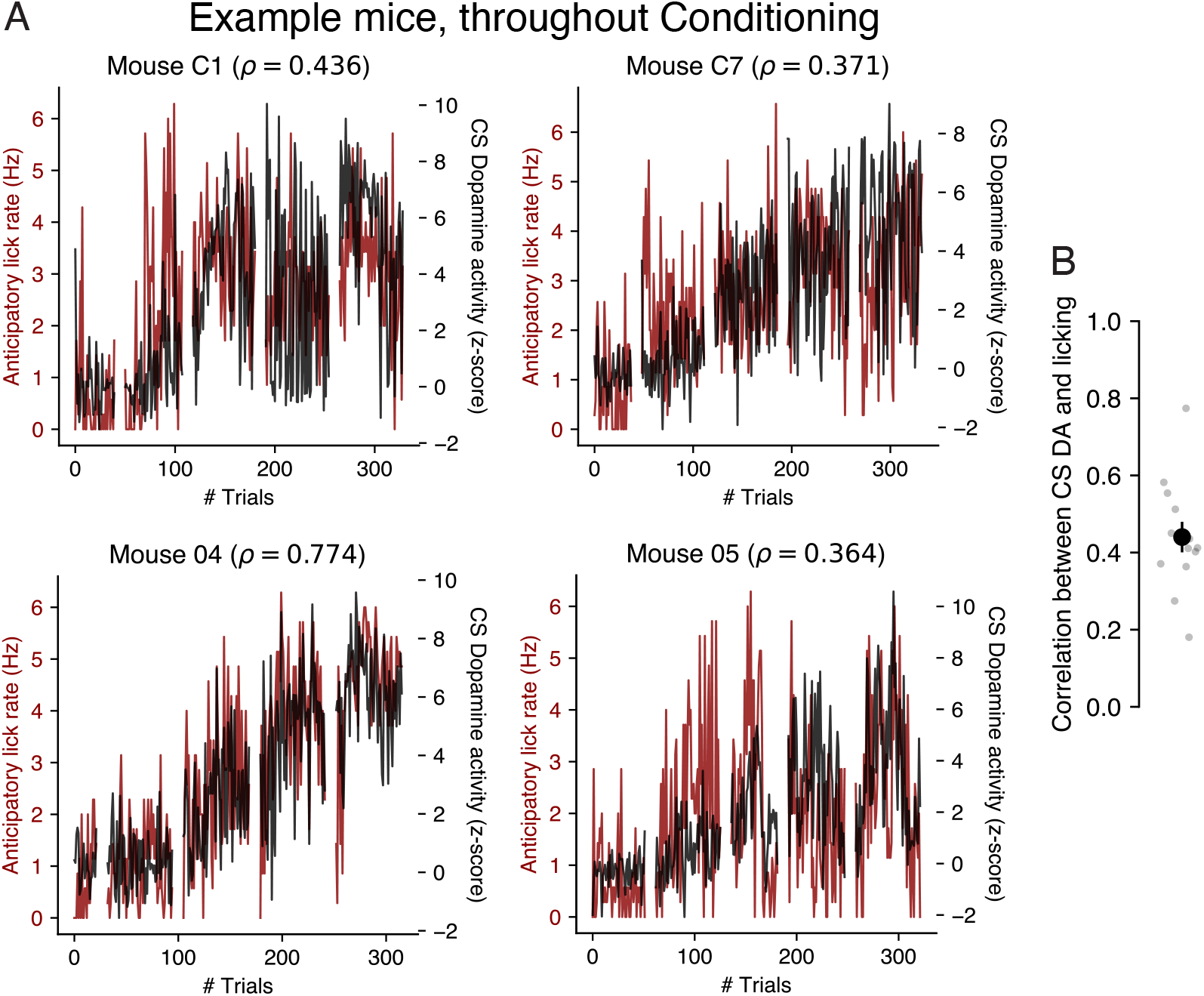
Trial-by-trial correlations between anticipatory lick rates and CS-evoked dopamine, throughout conditioning. **A**. Example time courses of anticipatory lick rate (red) and CS dopamine response (black) to the same cue on each trial of conditioning, with gaps of 10 trials added between sessions for visualization of session boundaries. Pearson correlation (*ρ*) between anticipatory lick rate and CS dopamine is shown above each subpanel. **B**. Pearson correlation between anticipatory lick rate and CS dopamine across conditioning trials for each mouse (grey dots), and across all mice (black circle). Black line depicts mean *±* SE.

Pearson correlation of 0.44*±*0.04 (mean *±* SE; *N* = 14 animals). While a TD learning agent’s estimate of CS value and its CS reward prediction error are expected to monotonically increase with learning, empirical measures of anticipatory licking and CS dopamine (assumed to reflect the CS reward prediction error) showed substantial non-monotonicity, suggesting this positive correlation may not be solely due to learning. Overall, our results suggest that trial-to-trial variability in CS dopamine can partially explain trial-to-trial variability in conditioned responding. As we will show next, this correspondence is *not* expected from a TD learning agent whose rates of conditioned responding reflect CS value.

### Anticipatory licking is maximally correlated with the immediately prior CS dopamine response

One caveat in interpreting the correlations between CS dopamine and licking above is that, even if licking were driven solely by value, TD learning would still predict correlations between the CS RPE (i.e., CS dopamine) and licking. This is because standard TD learning models predict that the RPE will gradually backpropagate across trials from the reward to the CS [4], as observed empirically [6]. This has been described as an ‘indirect’ relationship between the RPE/dopamine and responding [19], because the CS value on each trial will in general correlate with the RPEs from *previous* trials. Alternatively, the CS RPE may have a ‘direct’ (i.e., simultaneous) role in responding, in which case we would expect that the anticipatory response on trial *t* will depend maximally on the CS RPE on trial *t*. Thus, positive correlations between the CS RPE and anticipatory licking could, in principle, be explained both by the traditional assumption that licking reflects CS value, or by a model where licking is modulated directly by CS RPE.

To understand whether both of these hypotheses are indeed consistent with empirical data, we took a “phenotyping” approach (Fig 5A). First, we simulated thousands of TD learning agents, where each model had randomly sampled hyperparameters (see Methods). Next, we generated time series of anticipatory licking according to two different hypotheses. Under the *indirect* dopamine hypothesis (H1), licking is a readout of CS value. Under the *direct* dopamine hypothesis (H2), licking is a readout of the CS RPE/dopamine. For each resulting agent, we then assessed the Pearson correlation between the number of anticipatory licks on trial *t* and the magnitude of the CS RPE on trials *t − τ* for *τ* = 0, 1, …, 4. We refer to the set of resulting correlations, *β* ∈ ℝ^5^, as the agent’s *phenotype*.

**Fig 5.**
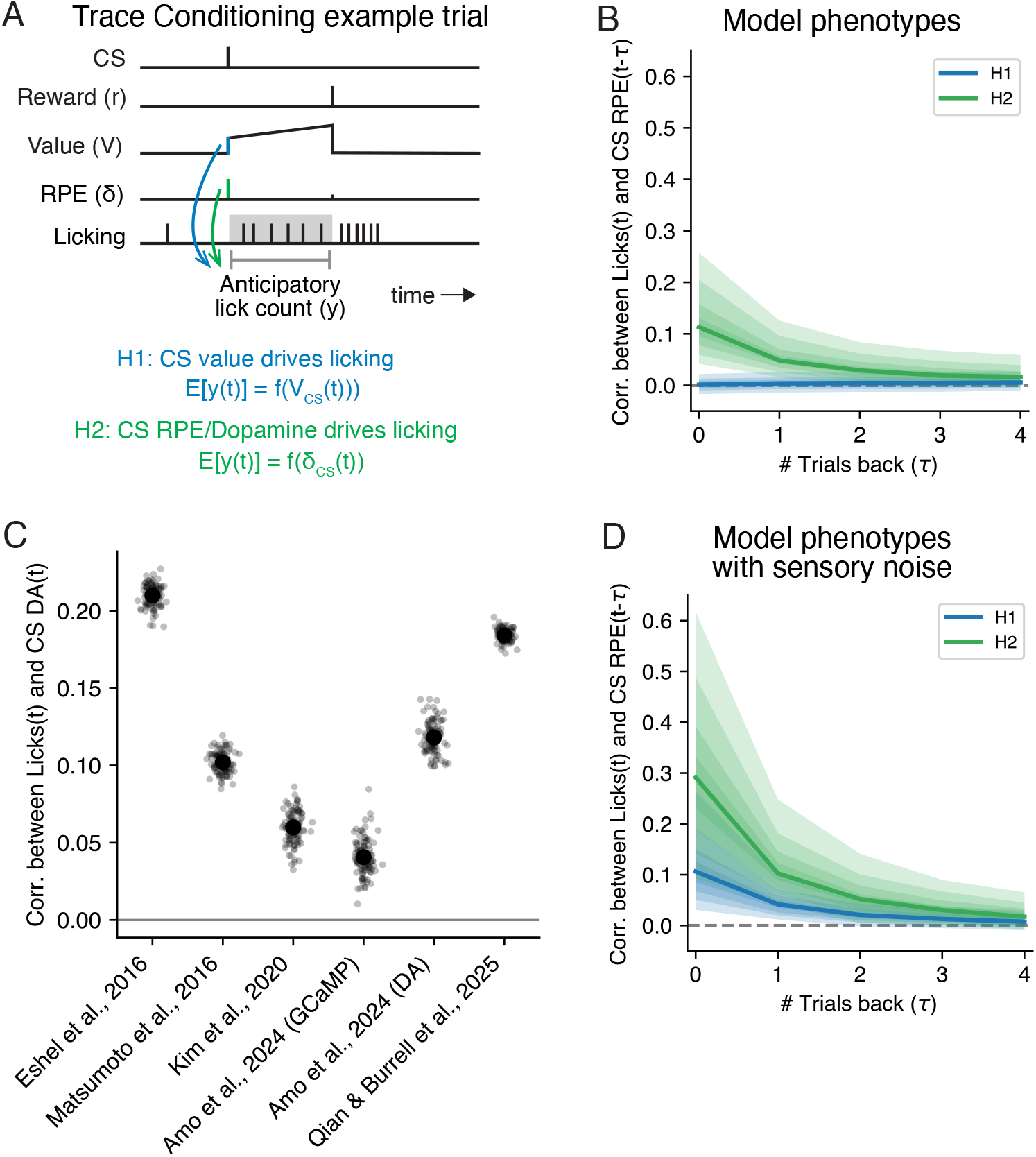
Phenotyping suggests anticipatory licking depends directly on CS RPEs/dopamine, or CS value with sensory noise. **A**. Schematic depicting two hypotheses for how the anticipatory lick count on a given trial, *y*(*t*), might depend on either CS value (*V*_*CS*_(*t*); H1) or on the CS RPE (*δ*_*CS*_(*t*); H2). **B**. Correlation between anticipatory lick count and CS RPE on previous trials, for simulated TD agents whose licking was generated according to H1 (blue) or H2 (green). Lines depict mean correlations across *N* = 500 models, and shading depicts deciles across models. **C**. Pearson correlations between anticipatory lick count and CS dopamine on the same trial, for empirical measurements from previous studies. Black dots indicate correlations across bootstraps, while black circles and lines indicate mean *±*SE across *N* = 1, 000 bootstraps. All correlations were statistically significant compared to correlations calculated using trial-shuffled DA. **D**. Same as panel B, but for agents with added sensory noise.

We found that the resulting phenotypes differed based on which agents had licking generated by CS value or by CS RPEs (Fig 5B). Agents whose licking directly reflected the CS RPE were distinguished by a positive correlation between anticipatory licking and the CS RPE on the same trial (i.e., *τ* = 0) (Fig 5B, green traces). Overall, agents whose licks reflected H2 versus H1 had larger correlations for *τ* = 0 (*d*^*′*^ = 1.751 *±* 0.077, mean *±* s.d. across *N* = 1, 000 bootstraps). Thus, this phenotyping approach could distinguish between two classes of TD agents where licking was driven either directly or indirectly by the RPE.

We next applied this same phenotyping approach to empirical data from all of the trace conditioning studies we considered earlier (see Methods). We found that anticipatory licking on a given trial was significantly positively correlated with the CS dopamine on the same trial across all previous studies (Fig 5C). This supports the idea that CS RPE/dopamine plays a direct role in driving conditioned responding during this experiment.

A positive correlation between CS dopamine and licking does not imply that the two are causally related. We reasoned that positive correlations could also arise if licking reflects CS value, but both CS value and licking have shared input noise. To implement this idea, we added noise to the CS sensory representation (e.g., reflecting a noisy sensory response to the odor) on each trial before applying TD learning, and generated licking according to the indirect model where licking reflects CS value. This addition of sensory noise produced similar, though weaker correlations between the CS RPE and licking on the same trial (Fig 5D). Nevertheless, the first regression weight was still consistently larger for agents whose licks reflected H2 versus H1 (*d*^*′*^ = 1.245 *±* 0.060, mean *±* s.d. across *N* = 1, 000 bootstraps).

Overall, our analyses revealed that, across a range of conditioning experiments, licking and CS dopamine were strongly positively correlated on the same trial, as expected by a TD model where CS RPEs (putatively signaled by dopamine) directly drive licking. Our phenotyping approach suggests that this correlation between CS dopamine and anticipatory licking on the same trial may be due either to a direct role for CS dopamine in licking, or to the presence of substantial noise in the CS sensory representation. Empirical support of the former possibility will be evaluated in a later section.

### Uncued peaks in CS dopamine between trials precede changes in licking

The analyses above revealed a correspondence between the concurrent CS RPE and conditioned responding. Our modeling suggests that this could be explained either by dopamine having a direct role in modulating licking, *or* a value-licking model with substantial noise in the sensory representation of the cue. One reason that these two possibilities are hard to distinguish is that both the CS RPE and CS value are directly dependent on the CS sensory representation. We reasoned that, if dopamine can directly modulate licking, then peaks in dopamine that are *not* cue-evoked may also yield changes in licking.

We therefore turned to comparing dopamine and licking during the intertrial interval (ITI) of data from [26], an epoch containing no experimentally-controlled inputs (Fig 6). Dopamine activity often showed substantial phasic activity during the ITI (Fig 6A), and lick rates were significantly larger in the 250 msec after versus before each peak (Fig 6B; Wilcoxon signed rank test *T* = 86.0, *p <* 1 *×* 10^*−*3^). We defined a “peak” in dopamine activity during the ITI as a time during the ITI at which the dopamine signal surpassed more than 3 standard deviations above average background levels across the entire experiment. We then aligned licking to each of these peaks and averaged across each peak, resulting in a “dopamine-peak-triggered average” of licking during the ITI. This analysis revealed an abrupt increase in licking following a dopamine peak during the conditioning phase (Fig 6C). This increase in licking was dose-dependent: the magnitude of these uncued dopamine peaks was positively correlated with larger subsequent increases in licking (Fig 6D). Importantly, because these dopamine peaks occurred during the ITI, they were uncued, and thus had zero objective value (because they were not predictive of reward). Nevertheless, these dopamine peaks preceded increases in licking, suggestive of a direct role for dopamine in modulating anticipatory licking.

**Fig 6.**
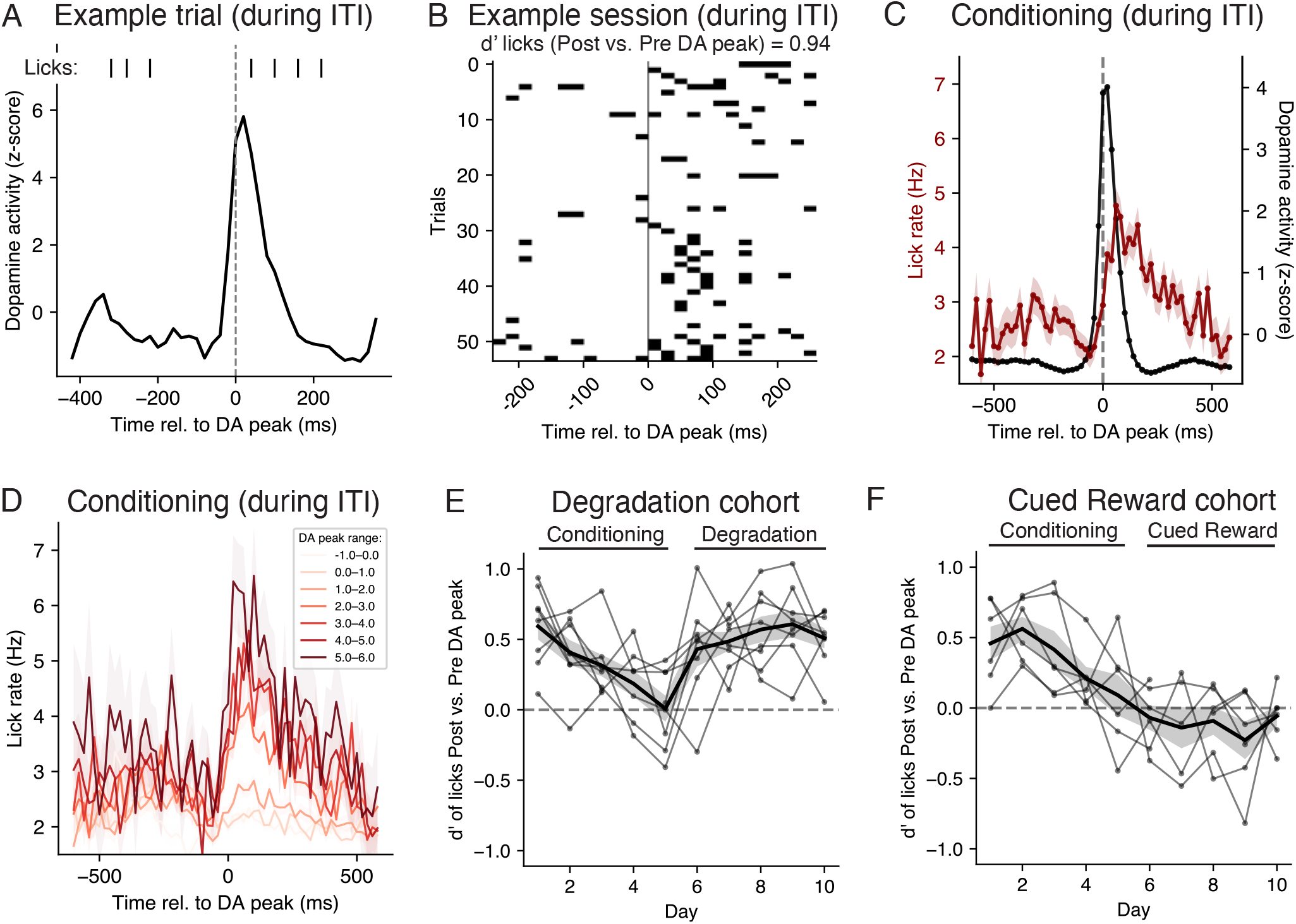
Uncued peaks in dopamine during intertrial intervals predict changes in lick rates. **A**. Licks and dopamine activity during the intertrial interval (ITI) of an example trial during a conditioning session, aligned to the time of a dopamine peak. **B**. Timing of licks relative to a dopamine peak during the ITI, for all ITIs from an example conditioning session with a dopamine peak. **C**. Average lick rate and dopamine activity relative to all ITI dopamine peaks from all conditioning sessions (mean *±*SE). **D**. Average lick rate (mean *±*SE) aligned to ITI dopamine peaks, grouped by different dopamine peak magnitudes. **E**. Sensitivity index (d’) in discriminating lick counts after versus before an ITI dopamine peak, for each mouse in the Degradation cohort on each session (black dots), and mean *±*SE across mice (black line and gray shading). **F**. Same as panel E, but for mice in the Cued Reward cohort.

To quantify this result for each session from each mouse, we compared the number of licks immediately before versus after the time of each dopamine peak, and summarized this difference using the sensitivity index (d’; see Methods). We then visualized the d’ of each cohort throughout the conditioning, degradation, and cued reward phases (Fig 6E-F). Across sessions of both cohorts, d’ was significantly positive throughout conditioning (Fig 6E-F; two-sided Wilcoxon signed rank test *T* = 160.0, *p <* 1*×*10^*−*3^), meaning the number of licks was consistently larger directly after versus before an uncued peak in dopamine during the ITI. For mice in the Contingency Degradation phase (i.e., some trials contained uncued rewards), d’ was also significantly positive (Fig 6E; two-sided Wilcoxon signed rank test *T* = 7.0, *p <* 1 *×* 10^*−*3^). However, for mice in the Cued Reward phase, d’ was no longer positive (Fig 6F; two-sided Wilcoxon signed rank test *T* = 98.0, *p* = 0.083). This result, where uncued peaks in dopamine preceded increases in licking during the ITI for some experimental conditions but not others, suggests a nonstationary or dynamic relationship between uncued peaks in dopamine and licking. While dopamine may directly modulate anticipatory licking, this relationship may be gated by other signals related to reward expectation, a possibility we will explore shortly.

### Causal manipulations of dopamine reveal direct roles for both CS RPE and value in driving conditioned responding

Whether or not the RPE has a direct role in driving conditioned responding can be evaluated empirically via causal perturbations of dopamine during or after Pavlovian conditioning. Previous perturbations include chronic dopamine depletion [34], and transient excitation or inhibition of midbrain dopamine neurons during the cue or reward periods [18, 15, 17]. Here we asked whether TD learning models can explain the results of these different dopamine perturbations. In particular, we were interested in whether explaining these results either ruled in or out the possibility of a direct role for CS RPEs/dopamine in driving conditioned responding.

We first considered results from a study that used optogenetics to transiently excite dopamine neurons during cue onset [17] (Fig 7A). The experimenters found that if dopamine neurons were stimulated during CS delivery, anticipatory licking was high for cues paired with reward (“CS+ Excitation”), but remained low for cues *not* paired with reward (“CS– Excitation”). This suggests that dopamine stimulation on its own was not sufficient to cause licking.

**Fig 7.**
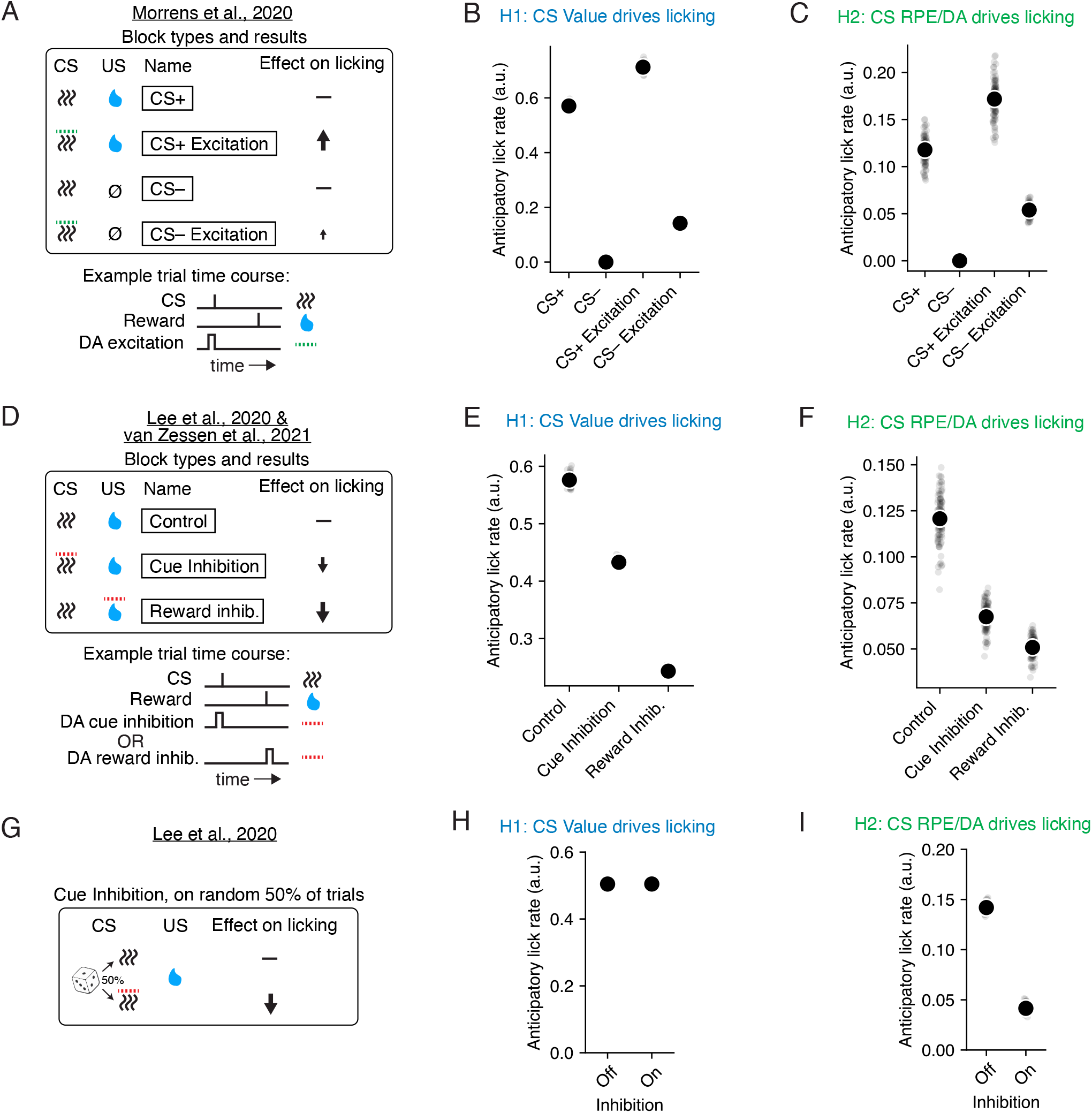
A model where the CS RPE drives anticipatory licking can explain the effects of causal manipulations of dopamine on licking. **A**. Top, four distinct conditions from [17], each with a distinct CS. “CS+” indicates a deterministically rewarded cue, while “CS–” indicates a deterministically unrewarded cue. For CS+ and CS– Excitation trials, optogenetic excitation of dopamine was modeled as a phasic, additive increase in the RPE during the CS delivery (see Methods). Anticipatory licking was high for CS+ Excitation and low for CS— Excitation. Bottom, example trial time course of a CS+ Excitation trial. **B**. Average lick rate for each trial type (each simulated separately), for models where lick rate is proportional to CS value (see Methods), across 100 experiments (black circles and lines indicate mean *±*SE for N=100 experiments). **C**. Same as panel B, but where lick rate is proportional to CS dopamine (see Methods). **D**. Top, three conditions and results from [15] and [18], each with a distinct CS. Optogenetic inhibition of dopamine was modeled as the opposite of excitation. Anticipatory licking was lowest in the Reward inhibition condition, but still reduced for Cue inhibition. Bottom, example trial time course of Cue inhibition versus Reward inhibition trials. **E-F**. Same conventions as panels B-C, but for the conditions in panel D. **G**. In a separate condition of [15], dopamine cue inhibition was either applied (“On”) or not applied (“Off”) during the delivery of a single CS on a random 50% of trials. Lick rates were reduced on the trials with cue inhibition. **H-I**. Same conventions as panels B-C, but for the conditions in panel G.

We applied TD learning models to these experiments, modeling dopamine excitation as a constant, *b*, added to the RPE, *δ*, simultaneous with the cue delivery (see Methods). We then simulated anticipatory licking as a readout of either CS value (Fig 7B) or CS RPE (Fig 7C). These are the same two models we considered earlier in Fig 5. While the results in [17] could be reproduced by TD learning models where CS value directly drove licking (Fig 7B), these results were also explained by a model where CS RPEs directly drove licking (Fig 7C). Notably, though the latter model has a direct link between the CS RPE and licking, pairing dopamine stimulation with cue presentation did *not* lead to large lick rates for the unrewarded CS– cue. This is because the CS RPE itself depends on CS value, which remains low for unrewarded cues. Thus, even TD models with a direct link between the CS RPE/dopamine and licking can nevertheless show a “gated” response to dopamine excitation. Because this result from [17] was explainable with both models of licking, these empirical results do not speak to whether or not the CS RPE has a direct role in anticipatory licking.

We next considered results from a pair of studies that used optogenetics to phasically *inhibit* dopamine activity during the cue or reward onset [15, 18] (Fig 7D). The authors found that dopamine inhibition during either the cue or the reward onset reduced anticipatory lick rates, with inhibition during the reward onset having a larger relative impact. We modeled dopamine inhibition as a constant, *b*, subtracted from the RPE/dopamine signal, *δ*, simultaneous with either the cue or reward delivery. We then applied TD learning models to these experiments as before (see Methods). We simulated anticipatory licking on each trial using either the CS value (Fig 7E) or CS RPE (Fig 7F) on each trial. Again we found that both models could reproduce the empirical results [15, 18], where lick rates were reduced for dopamine inhibition during either the cue or the reward onset, but with reward inhibition having a larger impact. Thus, these results do not speak to whether or not the CS RPE directly modulates anticipatory licking.

One reason why exciting or inhibiting dopamine can impact anticipatory licking even if CS RPE does not directly drive licking is due to the fact that these perturbations were applied in blocks of adjacent trials. These blockwise perturbations allow perturbed dopamine levels to impact CS value, and therefore licking, through learning. A more direct way of evaluating whether the CS RPE has an immediate influence on licking would be excite or inhibit dopamine activity on *random* trials rather than in blocks. Indeed, for TD models where licking was determined solely by CS value, randomly inhibiting CS RPE on random trials did *not* lead to reduced anticipatory licking compared to trials where CS dopamine was not inhibited (Fig 7G, top). By contrast, anticipatory licking was reduced during inhibited trials for TD models where licking was determined by CS RPE (Fig 7G, bottom). In fact, previous work did find that, when inhibiting dopamine during the cue period of random 50% of trials, anticipatory lick rates were lower on the randomly inhibited dopamine trials than on the uninhibited dopamine trials [15]. Thus, this empirical result combined with our simulations supports the idea that CS RPEs/dopamine can directly modulate anticipatory licking on the same trial, and that this modulation is distinct from the gradual changes in licking expected due to learning.

## Discussion

Using a combination of data analysis and computational modeling, we probed the relationship between phasic dopamine and anticipatory licking in mice during Pavlovian conditioning. Using data from multiple previously published studies, we found that the response of dopamine neurons to a reward-predictive cue (“CS dopamine”) was positively correlated with a higher anticipatory lick rate on the same trial. This suggests a link between the CS dopamine and the immediate rate of conditioned responding, a relationship not expected from the standard assumption that responding is proportional to CS value. In addition, we found that positing a direct link between the CS dopamine and anticipatory licking was necessary to reproduce previous empirical results showing that optogenetically inhibiting CS dopamine on random trials led to a decrease in anticipatory licking on those same trials. Taken as a whole, our results suggest that phasic dopamine directly modulates conditioned responding in addition to its canonical role as a learning signal. This direct role can be captured by a model in which the CS RPE drives conditioned responding.

The possibility that dopamine plays a role in action selection, and therefore conditioned responding, has been considered by previous computational models, mostly from the perspective of “incentive salience”— i.e., the role of dopamine in assigning motivational value to objects or actions [19, 22, 20, 35]. Previous studies using Pavlovian conditioning in rats have found that some rats develop “sign-tracking” behavior, where reward-predictive cues develop incentive motivational value, or incentive salience. This sign-tracking behavior is known to be dependent on dopamine [36, 20, 37, 38]. Incentive salience in tasks such as the ones we consider here, using odor cues, has not been studied to our knowledge. Previous work has found that dopamine release in the ventral striatum can initiate sniffing via action on both D1- and D2-expressing medium spiny neurons (MSNs) [39]. Sniffing could be one way of measuring sign-tracking of odor cues, in which case it would be interesting to evaluate the connection between trial-by-trial differences in sniffing, licking, and CS dopamine.

One potential circuit implementation consistent with a direct role for the CS RPE/dopamine in responding is if phasic dopamine provides feedforward excitation of striatal medium spiny neurons (MSNs). There is some empirical support of this idea [25], though this remains debated [40]. D1-MSNs are thought to modulate lick rate [41]. The response of MSNs to reward-predictive cues, putatively encoding CS value, may be modulated in a feedforward manner by CS dopamine, putatively encoding the CS RPE. Computational models based on this idea have been considered previously [42, 21].

Our findings add to a growing body of work on the role of phasic dopamine in movement vigor and initiation in other paradigms, spanning operant conditioning [42, 43, 44, 45], Pavlovian-instrumental transfer [46], and cue-potentiated feeding [27]. Closely related to our findings is work identifying a role for fast timescale changes in phasic dopamine activity in self-timed or self-initiated movements [47, 48, 49]. Our approach complements these studies by considering the relationship between dopamine and responding in the context of dopamine as the RPE signal in TD learning, which allows us to dissect the contributions of dopamine on responding that are immediate versus gradual through learning. An important direction for future work is to develop a unified computational framework that can explain the role of dopamine on both the timing and vigor of movements across these different paradigms.

Previous studies have considered the possibility that CS dopamine plays a direct role in modulating responding during Pavlovian conditioning. In [17], the authors dismissed this possibility because “CS dopamine stimulation in our experiments did not cause licking, per se, but selectively promoted responding only to rewarded cues.” As we showed in Fig 7A-C, however, this result on its own is not sufficient to discriminate between a TD-based model where licking is directly modulated by CS value or CS RPE. Fully discriminating between the roles of CS value and CS RPEs in modulating responding is challenging because, while we can measure putative RPEs using dopamine activity as a proxy, we cannot directly measure CS value. Another challenge is that causal perturbations to dopamine presented in the literature are typically applied in blocks rather than on randomly selected trials. This makes it challenging to distinguish between the learning and feedback-related effects of dopamine expected over the course of multiple trials, versus the immediate/feedforward effects expected on a single trial. The only study we are aware of that randomly perturbed dopamine activity during a Pavlovian trace conditioning task is [15], which we have shown here provides strong evidence that CS dopamine can directly modulate responding.

The slow correlations we observed between CS dopamine and conditioned responding in Fig 4 suggests that variability in an animal’s motivation or arousal may underlie some of the co-fluctuations in dopamine and licking. While arousal cannot explain the impacts of random optogenetic dopamine inhibition on licking (Fig 7G), arousal may nevertheless drive some co-fluctuations in both dopamine and licking. Previous work suggests a correspondence between conditioned responding and pupil size (an index for arousal) during trace conditioning tasks [50], but how and whether animals’ arousal differentially modulates CS dopamine and responding is an interesting, unanswered question.

The relationship between RPE and conditioned responding brings our work into contact with a broader set of ideas in animal learning theory [51, 52, 53, 54]. These accounts draw an important distinction between the number of training trials until a response acquisition criterion is met (sometimes called *associability*) and terminal response rate (after extensive training). It has been noted that reward expectation on its own is an inadequate predictor of associability. Rather, it is the *difference* between ISI (cue) and ITI (contextual) reward expectations that matters. The example of contingency degradation, discussed above, illustrates this point well: animals will not learn to produce conditioned responses to reward-predictive cues if the rate of reward is the same in the ISI and ITI [55]. This contrastive view of associability fits naturally with the contrastive nature of RPEs. However, here we have focused on the role of dopamine in driving terminal response rates, not associability, which we leave to future work.

In conclusion, our findings suggest that RPEs, putatively signaled by phasic dopamine activity, may directly modulate the intensity of conditioned responding on a trial-by-trial basis. This conclusion is supported by converging correlational evidence across multiple datasets, as well as the effects of random optogenetic stimulation of dopamine during cue presentation. Future work disentangling the contributions of sensory noise and arousal, as well as how dopamine could drive responding at the circuit level, will be important for building a more complete picture.

## Methods

### Analysis of dopamine activity and anticipatory licking

We analyzed previously collected dopamine and licking activity recorded from multiple trace conditioning studies conducted with mice [10, 32, 11, 33, 26]. For all studies, dopamine activity was analyzed in 50 msec bins. CS dopamine activity was defined using the activity within a 500 msec window following CS onset. For studies that recorded dopamine using electrophysiology [10, 32, 11], we used the average firing rate within the window. For studies that used fluorometry [33, 26], we used the maximum dopamine activity within the window. The anticipatory lick count on each trial was calculated as the number of licks during the ISI. Anticipatory lick rates were calculated as the anticipatory lick count divided by the ISI duration.

We describe below the trials we analyzed in each study.

- In [26], we analyzed dopamine activity measured using fluorometry with calcium sensors in ventral striatum (VS). We only analyzed trials to Odor A, the cue that was rewarded identically across all phases, with a reward probability of 75% and an ISI of 3.5 seconds.
- In [10], dopamine activity was measured using electrophysiology in the ventral tegmental area (VTA). We only analyzed trials to Odor C in the Variable-Expectation task only, which had a reward probability of 90% and an ISI of 2.0 seconds.
- In [32], dopamine activity was measured using electrophysiology in VTA. We only analyzed trials to Odor A in the high reward probability task, which had a reward probability of 90% and an ISI of 2.0 seconds.
- In [11], dopamine activity was measured using electrophysiology in VTA. We only analyzed trials to Odor B, which had a reward probability of 100% and an ISI of 1.5 seconds.
- In [33], dopamine activity was measured with fluorometry using calcium sensors (GCaMP) in VTA and dopamine sensors in VS. We analyzed these approaches separately. We only analyzed trials to Odor A, which had a reward probability of 80% and an ISI of 3.0 seconds.

### Uncued dopamine peaks

To identify uncued dopamine peaks (Fig 6), we identified time steps during the ITI where the z-scored dopamine activity surpassed 3 standard deviations of its activity. To ensure peaks were defined consistently for each mouse, z-scoring here involved normalizing across all sessions from the same mouse; results were similar when z-scoring per session. To summarize the discriminability of the distributions of lick counts before and after an uncued dopamine peak (Fig 6B,E,F), let the two distributions be indexed by *i* = 1 and *i* = 2, with mean *µ*_*i*_ and standard deviation *σ*_*i*_. Then we calculated the sensitivity index (d’) as *d*^*′*^ = (*µ*_2_ *− µ*_1_)*/σ*_1,2_, where 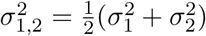 is the average of the variances.

### TD learning simulations

Here we will describe the TD learning procedure used in Fig 5 and Fig 7. Each simulated experiment consisted of concatenated trace conditioning trials for a single cue which was either deterministically rewarded (“CS+”) or unrewarded (“CS–”). The agent’s observations thus consist of the cue, *x*_*i*_ ∈ *{*0, 1*}*, and the reward, *r*_*i*_ ∈ *{*0, 1*}*, for all time steps *i* = 1, …, *J*. For simplicity, below we describe *x*_*i*_ and *r*_*i*_ for a single trial *t*, so that *J* is the total duration of this trial. The experiment itself is then made up of multiple concatenated trials.

Each trial had an intertrial interval (ITI) duration, and an interstimulus interval (ISI) duration. Each ITI was sampled from a Geometric distribution with parameter 0.25, plus a fixed duration of 5 time steps. Each ISI was a constant, *ISI* = 8.

In a given trial, *x*_*i*_ = 1 when *i* = 1 + *ITI*, and *x*_*i*_ = 0 otherwise. For a CS+ cue, *r*_*i*_ = 1 for *i* = 1 + *ITI* + *ISI*, and *r*_*i*_ = 0 otherwise. (For a CS– cue, *r*_*i*_ = 0 for all *i*.) Note that *J* = 1 + *ITI* + *ISI*.

To apply TD learning on trace conditioning trials, where the cue and reward delivery are separated in time, agents must have a sufficient representation of the cue. Following earlier work [1, 4], we used a complete serial compound representation, ***z***_*i*_ ∈ R^*K*^, where *K* = 15, with the first 6 entries corresponding to time steps within the ITI, the next 8 entries corresponding to time steps within the ISI, and the last entry serves as a bias term, i.e., *z*_*i*_(*K*) = 1 for all *i*. At each time step *i, z*_*i*_(*k*) = 1 for exactly one *k*, ignoring *k* = *K*. Thus, ***z***_*i*_ can be thought of as the agent’s representation of the current time step within a given trial. To model sensory noise, we added a noise term 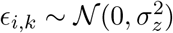 to each *z*_*i*_(*k*).

TD learning agents used linear value approximation: they learned a weight, ***w*** ∈ *ℝ*^*K*^, such that the agent’s value estimate at each time step was 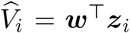. The agent updated ***w*** at each time step using the reward prediction error (RPE), *δ*_*i*_. To summarize, after concatenating all trials, TD learning then proceeded as follows, for all time steps *i* in the experiment:

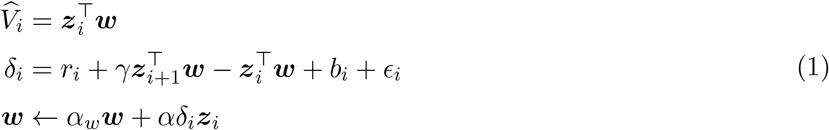

where *γ* = 0.9 is a discount factor, *b*_*i*_ = 0 except during dopamine perturbations (see below), *ϵ*_*i*_ *∼ 𝒩* (0, *σ*^2^) is i.i.d. Gaussian noise, *α*_*w*_ = 1 except when modeling value decay/forgetting, and *α* is the learning rate.

Note that in general we will use *i* to index time steps, and *t* to index trials. Also, 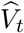 and *δ*_*t*_ will be assumed to be the value and RPE, respectively, at the time of CS onset during trial *t*.

### Phenotyping

To generate model phenotypes (Fig 5A,B,D), we first generated blocks of *N* = 10, 000 trace conditioning trials to a single deterministically rewarded cue. We then trained 500 TD learning agents for both H1 and H2, where each TD model used different hyperparameters (as defined in Eqn. (1)), sampled uniformly from the following options: *α* ∈ *{*0.001, 0.004, 0.01, 0.02, 0.05, 0.1*}, α*_*w*_ ∈ *{*1 *−* 10^*−*6^, 1 *−* 10^*−*5^, 1 *−* 10^*−*4^*}, σ* ∈ *{*0.1, 0.2, 0.3, 0.4*}*, and *α*_*y*_ ∈ *{*0.1, 0.2, …, 0.9*}*. Standard agents without sensory noise (Fig 5B) used *σ*_*z*_ = 0, while agents with sensory noise (Fig 5D) used *σ*_*z*_ = 0.1.

For each agent, the number of anticipatory licks on a given trial, *y*_*t*_, was given by:

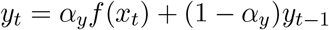

where 0 *< α*_*y*_ *≤* 1 induces exponential smoothing on *y*_*t*_, 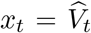 for agents in H1 (i.e., *x*_*t*_ is CS value), *x*_*t*_ = *δ*_*t*_ for agents in H2 (i.e., *x*_*t*_ is the CS RPE), and for simplicity the function *f* was set to the identity function, *f* (*x*) = *x*.

The phenotype of each agent consisted of the Pearson correlation between the number of anticipatory licks on each trial, *y*_*t*_, and the CS RPE on trial *t − τ, δ*_*t−τ*_, for each of *τ* = 0, 1, …, 4.

For empirical data (Fig 5C), we used the same phenotyping procedure, where *y*_*t*_ was the actual number of anticipatory licks on trial *t*, and *δ*_*t*_ was the CS dopamine activity. Lick counts and CS dopamine were combined across sessions from all animals after z-scoring per session to control for across-session variability, and padding with NaNs across sessions to ensure that correlations did not compare licking and CS dopamine from different sessions. Significance of correlations was assessed by comparing to the correlations found using trial-shuffled CS DA.

### Modeling dopamine perturbations

To model the effects of dopamine manipulations on anticipatory licking in Fig 7, for each trial type we generated 100 random experiments, each consisting of 1, 000 trace conditioning trials to a single cue which was either deterministically rewarded (“CS+”) or unrewarded (“CS–”). TD models (Eqn. (1)) were then fit to each of these experiments (separately for each cue) using learning rate *α* = 0.1, and sensory noise *σ*_*z*_ = 0.01. To model optogenetic excitation of dopamine for [17], we set *b*_*i*_ = 0.1 at the time of cue onset. To model optogenetic inhibition of dopamine for [15, 18], we set *b*_*i*_ = *−*0.1 either at the time of cue or reward onset, as indicated by the condition. For all other time steps we set *b*_*i*_ = 0. When modeling [17], perturbations were applied during all trials throughout learning, similar to the original experiments. When modeling [15, 18], perturbations were only applied after first training the TD models on 500 baseline trials. Each TD model was then used to generate anticipatory lick rates on each trial *t* based on either the CS value, *V*_*t*_, or CS RPE, *δ*_*t*_, using *p* = *f* (*V*_*t*_) or *p* = *f* (*δ*_*t*_). For simplicity, and to match what we did in the previous section regarding phenotyping, *f* was set to the identity function.

## Acknowledgements

This work was supported by the McNair Foundation (J.H.) and by NIH U19 NS113201 (N.U., S.J.G.).

## References

[1] Richard S Sutton and Andrew G Barto. Time-derivative models of pavlovian reinforcement. In Moore J Gabriel M, editor, Learning and computational neuroscience: Foundations of adaptive networks, pages 497–537. MIT Press, 1990.

[2] Elliot A Ludvig, Richard S Sutton, and E James Kehoe. Evaluating the td model of classical conditioning. Learning & behavior, 40(3):305–319, 2012.

[3] Samuel J Gershman. A unifying probabilistic view of associative learning. PLoS Computational Biology, 11(11):e1004567, 2015.

[4] Wolfram Schultz, Peter Dayan, and P Read Montague. A neural substrate of prediction and reward. Science, 275(5306):1593–1599, 1997.

[5] Jeremiah Y Cohen, Sebastian Haesler, Linh Vong, Bradford B Lowell, and Naoshige Uchida. Neuron-type-specific signals for reward and punishment in the ventral tegmental area. nature, 482(7383):85–88, 2012.

[6] Ryunosuke Amo, Sara Matias, Akihiro Yamanaka, Kenji F Tanaka, Naoshige Uchida, and Mitsuko Watabe-Uchida. A gradual temporal shift of dopamine responses mirrors the progression of temporal difference error in machine learning. Nature neuroscience, 25(8):1082–1092, 2022.

[7] Samuel J Gershman, John A Assad, Sandeep Robert Datta, Scott W Linderman, Bernardo L Sabatini, Naoshige Uchida, and Linda Wilbrecht. Explaining dopamine through prediction errors and beyond. Nature Neuroscience, 27(9):1645–1655, 2024.

[8] Hannah M Bayer and Paul W Glimcher. Midbrain dopamine neurons encode a quantitative reward prediction error signal. Neuron, 47(1):129–141, 2005.

[9] Neir Eshel, Michael Bukwich, Vinod Rao, Vivian Hemmelder, Ju Tian, and Naoshige Uchida. Arithmetic and local circuitry underlying dopamine prediction errors. Nature, 525(7568):243–246, 2015.

[10] Neir Eshel, Ju Tian, Michael Bukwich, and Naoshige Uchida. Dopamine neurons share common response function for reward prediction error. Nature neuroscience, 19(3):479–486, 2016.

[11] HyungGoo R Kim, Athar N Malik, John G Mikhael, Pol Bech, Iku Tsutsui-Kimura, Fangmiao Sun, Yajun Zhang, Yulong Li, Mitsuko Watabe-Uchida, Samuel J Gershman, et al. A unified framework for dopamine signals across timescales. Cell, 183(6):1600–1616, 2020.

[12] Elizabeth E Steinberg, Ronald Keiflin, Josiah R Boivin, Ilana B Witten, Karl Deisseroth, and Patricia H Janak. A causal link between prediction errors, dopamine neurons and learning. Nature Neuroscience, 16(7):966–973, 2013.

[13] Chun Yun Chang, Guillem R Esber, Yasmin Marrero-Garcia, Hau-Jie Yau, Antonello Bonci, and Geoffrey Schoenbaum. Brief optogenetic inhibition of dopamine neurons mimics endogenous negative reward prediction errors. Nature Neuroscience, 19(1):111–116, 2016.

[14] Benjamin T Saunders, Jocelyn M Richard, Elyssa B Margolis, and Patricia H Janak. Dopamine neurons create Pavlovian conditioned stimuli with circuit-defined motivational properties. Nature neuroscience, 21(8):1072–1083, 2018.

[15] Kwang Lee, Leslie D Claar, Ayaka Hachisuka, Konstantin I Bakhurin, Jacquelyn Nguyen, Jeremy M Trott, Jay L Gill, and Sotiris C Masmanidis. Temporally restricted dopaminergic control of reward-conditioned movements. Nature neuroscience, 23(2):209–216, 2020.

[16] Johann Du Hoffmann and Saleem M Nicola. Dopamine invigorates reward seeking by promoting cue-evoked excitation in the nucleus accumbens. Journal of Neuroscience, 34(43):14349–14364, 2014.

[17] Joachim Morrens, Çağatay Aydin, Aliza Janse van Rensburg, JoséEsquivelzeta Rabell, and Sebastian Haesler. Cue-evoked dopamine promotes conditioned responding during learning. Neuron, 106(1):142–153, 2020.

[18] Ruud Van Zessen, Jacques P Flores-Dourojeanni, Timon Eekel, Siem van den Reijen, Bart Lodder, Azar Omrani, Marten P Smidt, Geert MJ Ramakers, Geoffrey van der Plasse, Garret D Stuber, et al. Cue and reward evoked dopamine activity is necessary for maintaining learned pavlovian associations. Journal of Neuroscience, 41(23):5004–5014, 2021.

[19] David M Egelman, Christophe Person, and P Read Montague. A computational role for dopamine delivery in human decision-making. Journal of Cognitive Neuroscience, 10(5):623–630, 1998.

[20] Samuel M McClure, Nathaniel D Daw, and P Read Montague. A computational substrate for incentive salience. Trends in neurosciences, 26(8):423–428, 2003.

[21] Michael J Frank. Dynamic dopamine modulation in the basal ganglia: a neurocomputational account of cognitive deficits in medicated and nonmedicated parkinsonism. Journal of cognitive neuroscience, 17(1):51–72, 2005.

[22] Peter Dayan and Bernard W Balleine. Reward, motivation, and reinforcement learning. Neuron, 36(2):285–298, 2002.

[23] Rafal Bogacz. Dopamine role in learning and action inference. Elife, 9:e53262, 2020.

[24] Charles R Gerfen and D James Surmeier. Modulation of striatal projection systems by dopamine. Annual review of neuroscience, 34(1):441–466, 2011.

[25] Asha K Lahiri and Mark D Bevan. Dopaminergic transmission rapidly and persistently enhances excitability of d1 receptor-expressing striatal projection neurons. Neuron, 106(2):277–290, 2020.

[26] Lechen Qian, Mark Burrell, Jay A Hennig, Sara Matias, Venkatesh N Murthy, Samuel J Gershman, and Naoshige Uchida. Prospective contingency explains behavior and dopamine signals during associative learning. Nature neuroscience, pages 1–13, 2025.

[27] Andrew T Marshall, Briac Halbout, Angela T Liu, and Sean B Ostlund. Contributions of pavlovian incentive motivation to cue-potentiated feeding. Scientific reports, 8(1):2766, 2018.

[28] SD Paolo. Dopamine on d2-like receptors “reboosts” dopamine d1-like receptor-mediated behavioural activation in rats licking for sucrose. Neuropharmacology, 58(7):1085–1096, 2010.

[29] SD Paolo. Licking microstructure in response to novel rewards, reward devaluation and dopamine antagonists: possible role of d1 and d2 medium spiny neurons in the nucleus accumbens. Neuroscience & Biobehavioral Reviews, page 105861, 2024.

[30] Koji Toda, Nicholas A Lusk, Glenn DR Watson, Namsoo Kim, Dongye Lu, Haofang E Li, Warren H Meck, and Henry H Yin. Nigrotectal stimulation stops interval timing in mice. Current Biology, 27(24):3763–3770, 2017.

[31] Alam Coss, Ernesto Suaste, and Ranier Gutierrez. Lateral nac shell d1 and d2 neuronal ensembles concurrently predict licking behavior and categorize sucrose concentrations in a context-dependent manner. Neuroscience, 493:81–98, 2022.

[32] Hideyuki Matsumoto, Ju Tian, Naoshige Uchida, and Mitsuko Watabe-Uchida. Midbrain dopamine neurons signal aversion in a reward-context-dependent manner. Elife, 5:e17328, 2016.

[33] Ryunosuke Amo, Naoshige Uchida, and Mitsuko Watabe-Uchida. Glutamate inputs send prediction error of reward, but not negative value of aversive stimuli, to dopamine neurons. Neuron, 112(6):1001–1019, 2024.

[34] Martin Darvas, Amanda M Wunsch, Jeffrey T Gibbs, and Richard D Palmiter. Dopamine dependency for acquisition and performance of pavlovian conditioned response. Proceedings of the National Academy of Sciences, 111(7):2764–2769, 2014.

[35] Jun Zhang, Kent C Berridge, Amy J Tindell, Kyle S Smith, and J Wayne Aldridge. A neural computational model of incentive salience. PLoS Computational Biology, 5(7):e1000437, 2009.

[36] Kent C Berridge and Terry E Robinson. What is the role of dopamine in reward: hedonic impact, reward learning, or incentive salience? Brain Research Reviews, 28(3):309–369, 1998.

[37] Benjamin T Saunders and Terry E Robinson. The role of dopamine in the accumbens core in the expression of pavlovian-conditioned responses. European Journal of Neuroscience, 36(4):2521–2532, 2012.

[38] Amanda G Iglesias, Alvin S Chiu, Jason Wong, Paolo Campus, Fei Li, Jasmine K Bhatti, Shiv A Patel, Karl Deisseroth, Huda Akil, Christian R Burgess, et al. Inhibition of dopamine neurons prevents incentive value encoding of a reward cue: With revelations from deep phenotyping. Journal of Neuroscience, 43(44):7376–7392, 2023.

[39] Natalie L Johnson, Anamaria Cotelo-Larrea, Lucas A Stetzik, Umit M Akkaya, Zihao Zhang, Marie A Gadziola, Adrienn G Varga, Minghong Ma, and Daniel W Wesson. Dopaminergic signaling to ventral striatum neurons initiates sniffing behavior. Nature communications, 16(1):336, 2025.

[40] Charltien Long, Kwang Lee, Long Yang, Theresia Dafalias, Alexander K Wu, and Sotiris C Masmanidis. Constraints on the subsecond modulation of striatal dynamics by physiological dopamine signaling. Nature neuroscience, 27(10):1977–1986, 2024.

[41] Zhaorong Chen, Zhi-Yu Zhang, Wen Zhang, Taorong Xie, Yaping Li, Xiao-Hong Xu, and Haishan Yao. Direct and indirect pathway neurons in ventrolateral striatum differentially regulate licking movement and nigral responses. Cell reports, 37(3), 2021.

[42] Irene A Yun, Saleem M Nicola, and Howard L Fields. Contrasting effects of dopamine and glutamate receptor antagonist injection in the nucleus accumbens suggest a neural mechanism underlying cue-evoked goal-directed behavior. European Journal of Neuroscience, 20(1):249–263, 2004.

[43] Briac Halbout, Andrew T Marshall, Ali Azimi, Mimi Liljeholm, Stephen V Mahler, Kate M Wassum, and Sean B Ostlund. Mesolimbic dopamine projections mediate cue-motivated reward seeking but not reward retrieval in rats. Elife, 8:e43551, 2019.

[44] Sarah Fischbach-Weiss, Rebecca M Reese, and Patricia H Janak. Inhibiting mesolimbic dopamine neurons reduces the initiation and maintenance of instrumental responding. Neuroscience, 372:306–315, 2018.

[45] Abigail Kalmbach, Vanessa Winiger, Nuri Jeong, Arun Asok, Charles R Gallistel, Peter D Balsam, and Eleanor H Simpson. Dopamine encodes real-time reward availability and transitions between reward availability states on different timescales. Nature communications, 13(1):3805, 2022.

[46] Anja Lex and Wolfgang Hauber. Dopamine d1 and d2 receptors in the nucleus accumbens core and shell mediate pavlovian-instrumental transfer. Learning & memory, 15(7):483–491, 2008.

[47] Joaquim Alves Da Silva, Fatuel Tecuapetla, Vitor Paixão, and Rui M Costa. Dopamine neuron activity before action initiation gates and invigorates future movements. Nature, 554(7691):244–248, 2018.

[48] Allison E Hamilos, Giulia Spedicato, Ye Hong, Fangmiao Sun, Yulong Li, and John A Assad. Slowly evolving dopaminergic activity modulates the moment-to-moment probability of reward-related self-timed movements. Elife, 10:e62583, 2021.

[49] Allison E Hamilos, Isabella C Wijsman, Qinxin Ding, Pichamon Assawaphadungsit, Zeynep Ozcan, Elias Norri, Kimberly Reinhold, Bernardo L. Sabatini, and John A Assad. Dopamine reward transients calibrate movement timing via one-shot updates to behavioral vigor. bioRxiv, 2025.

[50] Laurens Winkelmeier, Carla Filosa, Renée Hartig, Max Scheller, Markus Sack, Jonathan R Reinwald, Robert Becker, David Wolf, Martin Fungisai Gerchen, Alexander Sartorius, et al. Striatal hub of dynamic and stabilized prediction coding in forebrain networks for olfactory reinforcement learning. Nature Communications, 13(1):3305, 2022.

[51] CR Gallistel and John Gibbon. Time, rate, and conditioning. Psychological Review, 107(2):289–344, 2000.

[52] Charles Randy Gallistel. Reconceptualized associative learning. Perspectives on Behavior Science, 48(2):203–239, 2025.

[53] Samuel J Gershman. Bridging computation and representation in associative learning. Computational Brain & Behavior, pages 1–15, 2025.

[54] Justin A Harris and Charles Randy Gallistel. Information, certainty, and learning. eLife, 13:RP102155, 2026.

[55] Robert A Rescorla. Probability of shock in the presence and absence of CS in fear conditioning. Journal of Comparative and Physiological Psychology, 66(1):1, 1968.

